# A circadian-like gene network regulates heterochronic miRNA transcription in *C. elegans*

**DOI:** 10.1101/2022.09.26.509508

**Authors:** Brian Kinney, Shubham Sahu, Natalia Stec, Kelly Hills-Muckey, Dexter W. Adams, Jing Wang, Matt Jaremko, Leemor Joshua-Tor, Wolfgang Keil, Christopher M. Hammell

**Author notes:** Correspondence (W.K.); (C.M.H.) (lead contact). These authors contributed equally (Brian Kinney, Shubham Sahu).

## Abstract

Developmental robustness relies on precise control of the timing and order of cellular events. In *C. elegans*, the invariant sequence of post-embryonic cell fate specification is controlled by oscillatory patterns of heterochronic microRNA transcription that are phase-locked with the larval molting cycle^1-4^. How these transcriptional patterns are generated and how microRNA dosage is controlled is unknown. Here we show that transcriptional pulses of the *lin-4* heterochronic microRNA are produced by two nuclear hormone receptors, NHR-85 and NHR-23, whose mammalian orthologs, Rev-Erb and ROR, function in the circadian clock. While Rev-Erb and ROR play antagonistic roles in regulating once-daily transcription^5-7^, we find that NHR-85 and NHR-23 bind cooperatively as heterodimers to *lin-4* regulatory elements to induce a single brief pulse of expression during each larval stage. We demonstrate that the timing and duration of *lin-4* transcriptional pulses are programmed by the phased overlap of NHR-85 and NHR-23 protein expression and that these regulatory interactions are post-transcriptionally controlled by LIN-42, the circadian Period ortholog in *C. elegans*. These findings suggest that an evolutionary rewiring of the circadian clock machinery is co-opted in nematodes to generate periodic transcriptional patterns that define cell fate progression.

## Introduction

Many gene regulatory networks encode and read information controlling the tempo and order of developmental events. *C. elegans* postembryonic maturation is compartmentalized into four larval stages with distinct patterns of cell division, cell differentiation, and cuticle formation that are separated by molts^8^. The transition from one stage-specific pattern of cell division to the next is mediated by the accumulation of heterochronic microRNAs (miRNAs) that post-transcriptionally downregulate temporal identity target genes that define stage-specific gene expression patterns^9^. Hence, lower or higher dosages of heterochronic miRNAs result in precocious cellular transitions or reiteration of stage-specific cellular programs, respectively. The transcription of heterochronic microRNAs is periodic, peaking at specific phases of each larval molting cycle^1-4^. This suggests the existence of a clock-like system to coordinate their temporal expression with overall developmental progression.

Evidence suggests that LIN-42, the *C. elegans* ortholog of the circadian Period protein, is required for temporal patterning by directly modulating the dynamic features of heterochronic microRNA transcription^1,2,4^. Loss-of-function mutations in *lin-42* increase the amplitude and duration of cyclical heterochronic miRNA transcription, indicating that LIN-42, like its mammalian ortholog, functions as a transcriptional repressor^1,2,4^. Consequently, *lin-42* mutants develop precociously due to overaccumulation of heterochronic microRNAs^2,10^. While most core components of the heterochronic gene regulatory network are expressed in a graded temporal pattern, LIN-42 is highly dynamic, with a single peak of expression during each larval stage^2,10^. The similar molecular functions and expression dynamics between LIN-42 and mammalian Period suggest that a regulatory architecture akin to the one that generates the once-daily circadian transcriptional patterns in mammals may play a role in temporal cell fate specification in nematodes. Intriguingly, *C. elegans* lacks orthologs of CLOCK and BMAL1, the central transcription factors that drive oscillatory circadian transcription and the direct targets of Period repression. Thus, the mechanism by which LIN-42 modulates the transcriptional output of *C. elegans* heterochronic microRNAs is currently unknown.

### Oscillatory *lin-4* transcription is pulsatile

Previous measurements of miRNA transcription used destabilized GFP reporters that must be transcribed, processed, and translated to visualize expression dynamics^2,11^. These features limit their temporal resolution. We used the MS2/MCP-GFP tethering system, where engineered RNA loops derived from MS2 bacteriophage and a co-expressed <u>M</u>S2 <u>c</u>oat <u>p</u>rotein fused to green fluorescent protein (MCP-GFP) can be concentrated and localized at a gene of interest by active transcription (Fig.1a)^12^. We measured the transcriptional dynamics of the *lin-4* heterochronic miRNA that is expressed periodically throughout all larval stages and down-regulates *lin-14* expression within the heterochronic gene regulatory network early in development^11^. We generated a transgene harboring 24 copies of a synthetic MS2 hairpin immediately downstream of the *lin-4* pre-miRNA (Fig. 1a and Extended Data Fig. 1a)^11,13^. This transgene rescues *lin-4* null allele phenotypes (Extended Data Fig. 1b,c). We also ubiquitously expressed MCP-GFP to detect MS2-tagged RNAs and a histone::mCherry fusion to locate nuclei (Extended Data Fig. 1a). Examination of transgenic animals revealed that the formation of nuclear MCP-GFP foci occurred at each larval stage in somatic tissue types known to transcribe *lin-4* (Fig. 1b). MCP-GFP foci were not observed in developing embryos (n > 50) or in starvation arrested L1 larva (n = 23); consistent with the activation of *lin-4* transcription after the initiation of larval development^14^.

**Fig. 1.**
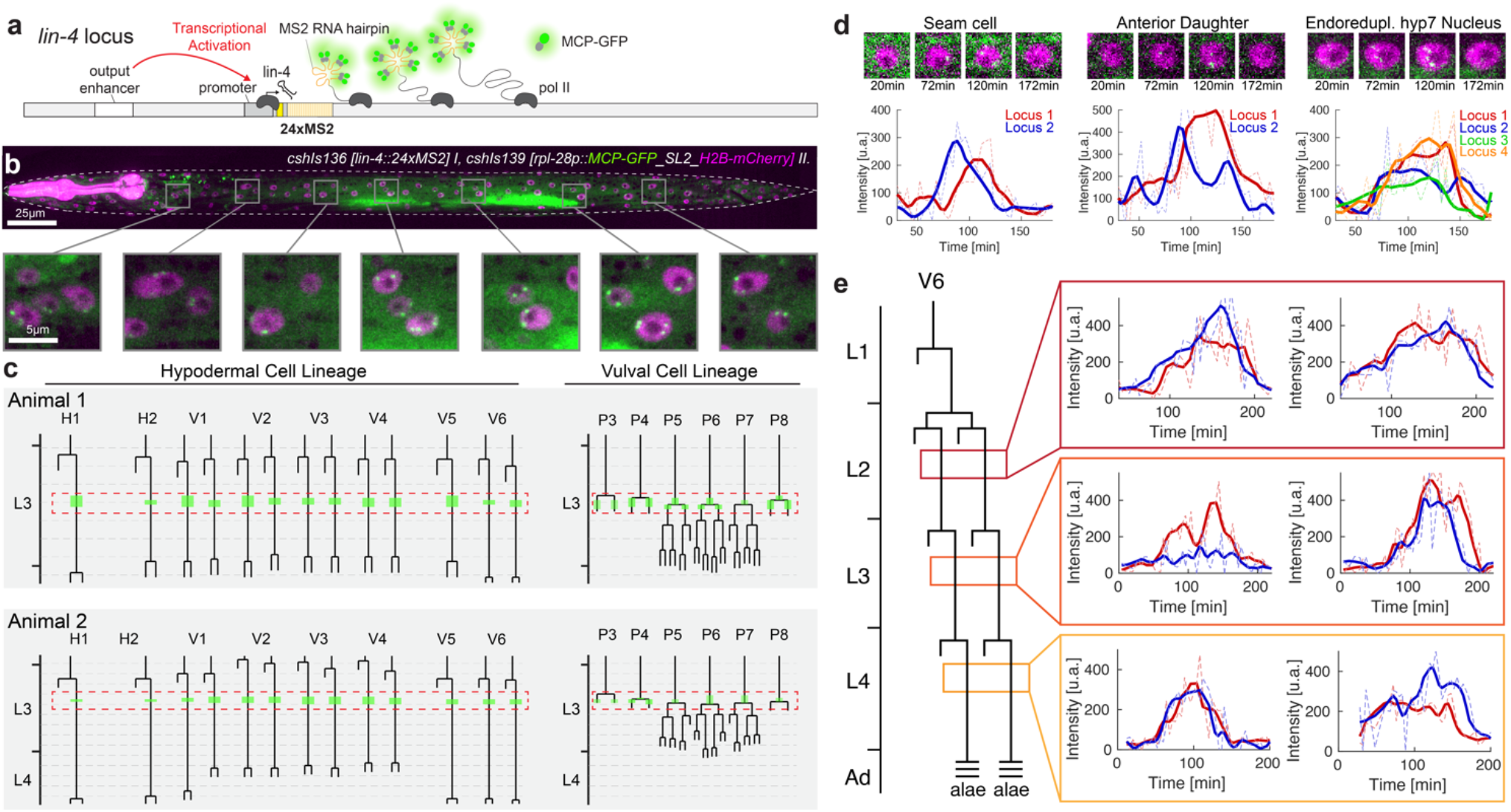
*lin-4* transcription is highly pulsatile at each larval stage. (**a**) The MS2/MCP system is composed of an MS2 coat protein GFP fusion (MCP-GFP) which can bind to MS2 RNA hairpins engineered into primary miRNA transcripts. (**b**) Image of mid-L3 staged larva and magnified insets showing MCP-GFP spots in hypodermal cells along the anteroposterior axis. (**c**) Two examples of L3/L4 hypodermal and vulval precursor cell lineages, with overlayed expression patterns indicating when MCP-GFP foci were visible in each lineage. Red boxes show the temporal region where coordinated *lin-4* transcription is observed. Dashed horizontal grey bars indicate hours. (**d**) Snapshots of individual seam cell and hyp7 cell expression trajectories from L3-staged animals. (**e**) Expression traces in pairs of V6.p seam cells from L2, L3, and L4 staged animals.

To directly determine the level of coordination of transcriptional programs, at single-cell resolution, across cell and tissue types in living animals, we used our microfluidics-based platform^15^ for long-term imaging of *C. elegans* larvae harboring *lin-4::24xMS2/MCP-GFP* system. We quantified expression dynamics in hypodermal cells where the timing of transcriptional activity can be accurately assessed in relation to the occurrence of stage-specific cell division patterns^8^. We screened for periods of *lin-4* transcriptional activity by imaging at 15min time intervals from the first larval stage (L1) to the mid-L4 stage (∼60h) (n>10). *lin-4::24xMS2* transcription was highly pulsatile, with a single transcriptional episode of ∼45-90min in each cell during each larval stage, followed by long periods of inactivity (>8hrs at 20°C) (Fig. 1c). Transcriptional activation across cells within the hypodermis was highly concordant, exhibiting similar transcriptional on and off times for all *lin-4::24xMS2* loci (Fig. 1e). For instance, during the mid-L3 stage, the appearance of *lin-4::24xMS2* expression throughout the hypodermis generally occurred within minutes of the first vulval precursor cell (VPC) divisions (P3.p or P4.p) and were completed by the divisions of remaining VPCs (P5.p-P7.p) (n = 15) (Fig. 1c).

To determine how similar the transcriptional dynamics in individual cells within the hypodermis are, we performed short-term imaging time courses (<6h) at 4min intervals in staged larvae. During transcriptional episodes, we detected near synchronous accumulation of MCP-GFP foci at each hypodermal *lin-4::24xMS2* locus for 60-90 minutes (Fig. 1d,e, Suppl. Movie 1,2) (>15 animals). We found no signs of “bursty” transcription^16^ as MCP-GFP foci were continuously maintained at each *lin-4::24xMS2* transgene for the entire transcriptional episode (Suppl. Movie 1,2). These features were independent of the number of *lin-4::24xMS2* loci per nucleus as cell types that undergo endoreduplication (i.e., hyp7 cells) exhibited MCP-GFP foci dynamics indistinguishable from diploid cells (Fig. 1d). The dynamic features of *lin-4::24xMS2* expression in hypodermal cells were similar across different developmental stages (Fig. 1e) suggesting that the same regulatory programs controlling *lin-4* transcription were repeated at each larval stage. Pulsatile transcription also occurs in a broad array of additional cell types that normally express *lin-4* (including the pharynx and intestinal cells) (Extended Data Fig. 2). Therefore, the gene regulatory network that generates *lin-4* transcriptional pulses at each stage of development organizes the timing, amplitude and duration of transcription in a highly reproducible manner.

### NHR-85 and NHR-23 function cooperatively

Full transcriptional activation of *lin-4* requires a conserved upstream regulatory element, the *<u>P</u>*ulse *<u>C</u>*ontrol *<u>E</u>*lement (PCE), located ∼2.8kb upstream of the *lin-4* sequence ^11^. We performed a yeast-one-hybrid screen to discover TFs that bind this regulatory region using the entire 514bp PCE as bait. We identified three TFs that specifically bound the PCE (BLMP-1, NHR-23, and NHR-85) (Fig. 2a). BLMP-1 controls the amplitude of *lin-4* expression and functions as a pioneer factor to decompact the *lin-4* locus throughout development ^11^. *nhr-85* and *nhr-23* encode two nuclear hormone receptors (NHRs) that are the closest nematode orthologs of human circadian TFs Rev-Erb and ROR, respectively (Fig. 2b). NHR-23 expression cycles with the larval molts and has been implicated in controlling the pace of larval development ^17-19^. Furthermore, *nhr-23* and the terminal heterochronic miRNA, *let-7*, genetically interact to mutually limit supernumerary molts during adulthood ^18,20^. Analysis of publicly available ChIP-seq data indicated that all three TFs interact *in vivo* with *lin-4* regulatory sequences (Fig. 2a), and their binding sites are enriched in the promoters of cyclically expressed mRNAs and other heterochronic miRNAs (Extended Data Fig. 3) (table S1 and S2).

**Fig. 2.**
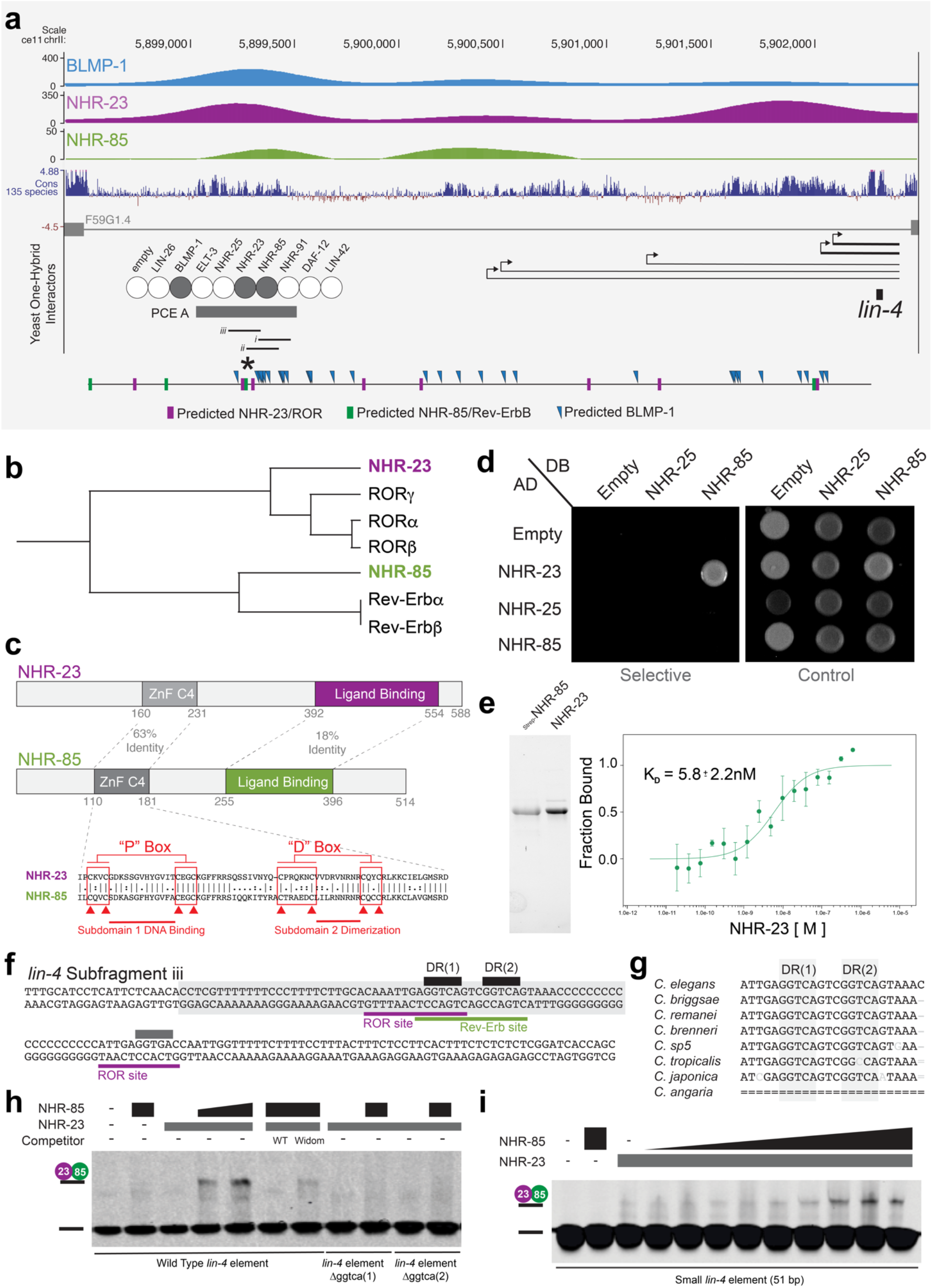
NHR-85^Rev-Erb^ and NHR-23^ROR^ form a heterodimeric complex that binds cooperatively to regulatory elements controlling pulsatile *lin-4* transcription. (**a**) BLMP-1, NHR-85, and NHR-23 bind to the *lin-4* PCE in one-hybrid assays^11^. Browser tracks showing BLMP-1, NHR-23, and NHR-85 bindings sites near the *lin-4* locus. Also indicated are the major *lin-4* pri-miRNAs^13^, the computationally defined binding sites for each TF, and the sub-fragments of the *lin-4* PCE element used in gel shifts below. The asterisk indicates the location of the direct repeats of GGTCA in the PCE element. (**b**) Sequence relationships between NHR-23 and NHR-85 and human Rev-Erb and ROR. (**c**) NHR-23^ROR^ and NHR-85^Rev-Erb^ domain organization. (**d**) NHR-23^ROR^ and NHR-85^Rev-Erb^ interact with each other in two-hybrid assays. (**e**) Recombinant, purified strep-NHR-85^Rev-Erb,^ and NHR-23^ROR^ interact in MST binding assays. (**f**) Sequences of the PCE sub-fragment iii from a and the 51bp minimal binding element (grey box) derived from *lin-4* PCEiii. (**g**) Conservation of the direct repeats DRs in the PCE elements of different nematode species. (**h** and **i**) EMSA experiments of wild-type and mutant target DNAs using recombinant NHR-85^Rev-Erb^ and NHR-23^ROR^.

Nuclear hormone receptors often bind cooperatively as homo- or hetero-dimeric complexes at closely spaced cis-regulatory DNA elements ^21-23^. Several features of NHR-85^Rev-Erb^, NHR-23^ROR,^ and the *lin-4* PCE suggest that this may also be the case for *lin-4* transcription. First, NHR-85^Rev-Erb^ and NHR-23^ROR^ share significant sequence homology within their C4-type Zinc finger DNA-binding domains, suggesting they bind similar sites (Fig. 2c) ^24^. Second, we found that NHR-85^Rev-Erb^ and NHR-23^ROR^ heterodimerize in yeast two-hybrid assays and *in vitro* with high affinity (5.8 +/- 2.2 nM K_D_ (Fig. 2d,e)). Third, sequences within the PCE element contain direct GGTCA repeats that are predicted binding sites of NHR-85^Rev-Erb^ and NHR-23^ROR^ (Fig. 2f and g). Fourth, we found that concentrations of NHR-23^ROR^ that were insufficient to bind the PCE alone were dramatically stimulated by the addition of NHR-85^Rev-Erb^, indicating cooperative binding (Fig. 2h, i). Supporting this, NHR-85^Rev-Erb^ alone could not bind *lin-4* PCE DNA fragments (Extended Data Fig. 4c). Finally, the cooperative binding of NHR-85^Rev-Erb^ and NHR-23^ROR^ required both GGTCA repeats, suggesting they bind closely spaced regulatory elements as a heterodimer (Fig. 2h).

### NHR-85/NHR-23 expression coincides with *lin-4* transcription

We quantified mRNA and protein expression during post-embryonic development to determine how NHR-85, NHR-23, and LIN-42 expression patterns may contribute to the regulation of oscillatory lin-4 transcription. Expression of *nhr-85* begins from an L1-stage arrest with a pulse of transcription that is followed by a monotonic expression pattern for the remaining larval stages (Fig. 3a). In contrast, *nhr-23* and *lin-42* mRNAs were expressed in phased, high-amplitude oscillatory patterns (Fig. 3a). We next explored the temporal dynamics of the corresponding proteins, by quantifying the expression of endogenously-tagged alleles during the L4 stage, where changes in vulval morphogenesis can be directly correlated with developmental age ^25^. NHR-23^ROR^::mScarlet and LIN-42^Period^::GFP are dynamically expressed in all hypodermal cells, with a single peak of expression that matched the phased expression of their mRNAs (Fig. 3c). We also found that NHR-85^Rev-Erb^::GFP expression was highly dynamic during these periods, indicating substantial post-transcriptional regulation of expression in the L2-L4 stages of development. Specifically, peak expression of NHR-85^Rev-Erb^::GFP occurred at ecdysis (shortly before NHR-23^ROR^::mScarlet onset), becomes undetectable by the L4.3 stage of development, and resumes expression at the L4.6 stage in an antiphasic manner to the expression pattern of LIN-42^Period^::YFP (Fig. 3b). Highly similar phased expression patterns of these proteins are also maintained in L4-staged vulval tissues (Fig. 3c) as well as in L3-staged hypodermal cells and VPCs (Fig. 3c-e).

Since NHR-85^Rev-Erb^ and NHR-23^ROR^ heterodimerize and bind cooperatively to regions of the *lin-4* enhancer that control dynamic transcription, we hypothesized that the 60-90min pulses of *lin-4* transcription might only occur in the short window of each larval stage where NHR-85^Rev-Erb^ and NHR-23^ROR^ are co-expressed. To compare the timing of these events, we examined MCP-GFP localization in vulval cells during the L3 and L4 stages, where the rapid, stereotyped vulval cell division patterns^15^ and changes in morphology^25^ enable precise determination of the timing of *lin-4::24xMS2* transcription and TF expression dynamics. We found a correspondence between NHR-85^Rev-Erb^ and NHR-23^ROR^ co-expression and *lin-4::24xMS2* transcription in both vulval and hypodermal cells (Fig. 3d,e). While the expression of both nuclear receptors is phased, the transient expression of the *lin-4::24xMS2* transgene only occurs during the brief period when both NHRs are expressed. The timing of NHR-85^Rev-Erb^ post-transcriptional downregulation is also concurrent with the onset of LIN-42^Period^ expression in VPCs and hypodermal cells (Fig. 3c-e). This suggests that the dynamic patterns of these TFs control the timing and duration of *lin-4* transcriptional pulses.

**Fig. 3.**
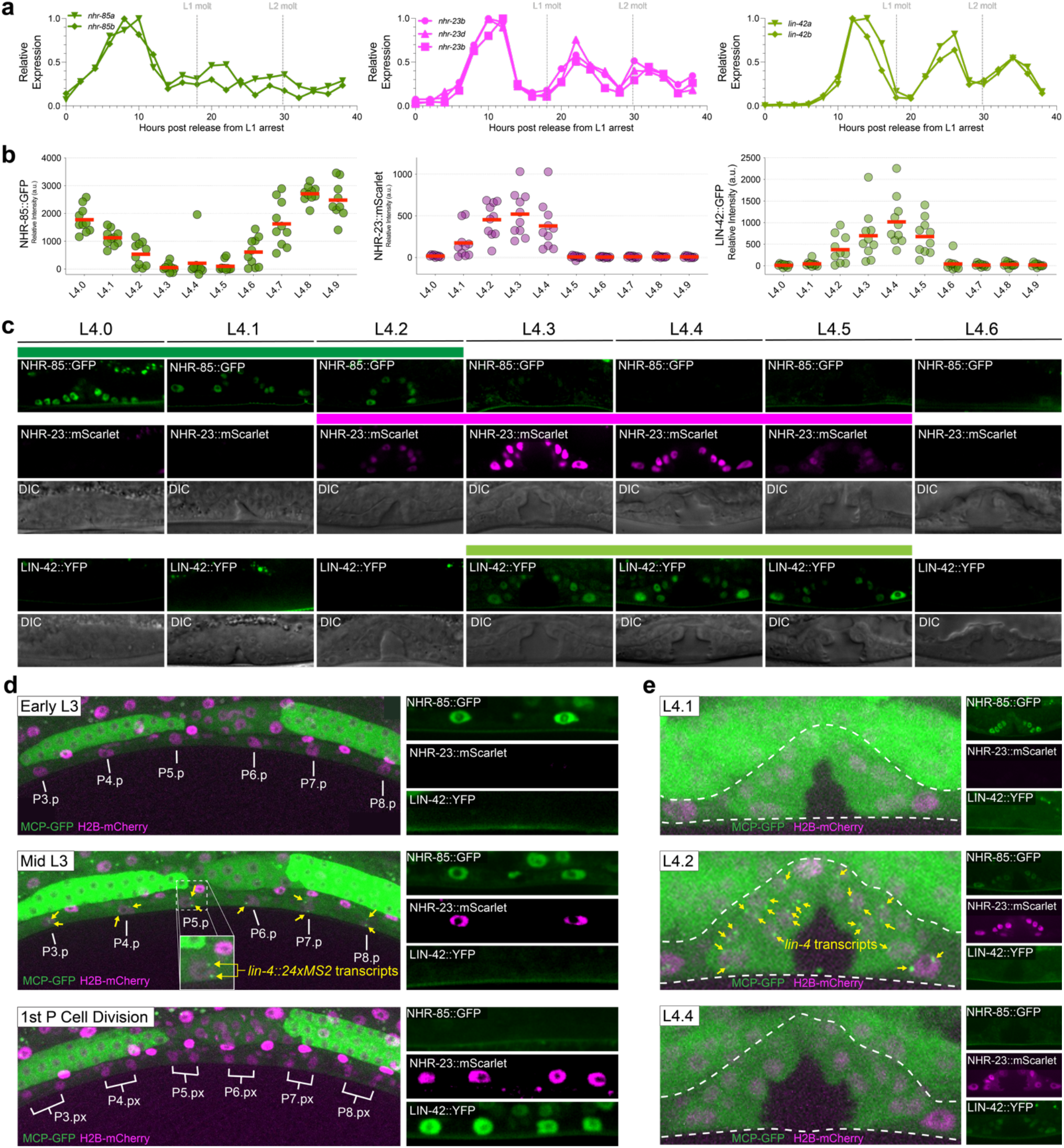
*lin-4* transcription occurs during the brief periods where NHR-85^Rev-Erb^::GFP and NHR-23^ROR^::mScarlet are co-expressed. (**a**) RNA-seq time course data of *nhr-85, nhr-23*, and *lin-42* mRNA expression patterns^3^. (**b-c**) Quantification and micrographs depicting NHR-85^Rev-Erb^::GFP, NHR-23^ROR^::mScarlet, and LIN-42^Period^::YFP expression in hypodermal seam cells and vulval cells, respectively, in each morphologically defined L4 substage^25^. Circles in b represent average measurements from individual animals (3 cells sampled); red bars indicate the mean. Colored bars indicate ranges of detectable expression. (**d**) Time course experiments demonstrate that *lin-4::24xMS2* expression occurs immediately before the first Pn.p cell divisions and not until NHR-85^Rev-Erb^::GFP are NHR-23^ROR^::mScarlet are co-expressed in the VPCs. *lin-4::24xMS2* expression terminates by the time NHR-85^Rev-Erb^::GFP expression is extinguished around the time of the first VPC division. (**e**) Dynamic *lin-4::24xMS2* transcription also correlates with NHR-85^Rev-Erb^::GFP/ NHR-23^ROR^::mScarlet in the L4 stages of vulval development.

### LIN-42^Period^ posttranscriptionally represses NHR-85

Human Per2 (PERIOD2) protein interacts with multiple mammalian NHRs (including Rev-Erb) to modulate their transcriptional activity ^26^. To test whether LIN-42^Period^ physically interacts with either NHR-85^Rev-Erb^ or NHR-23^ROR^, we used two-hybrid assays ^27^. We found that both major LIN-42 isoforms interact with NHR-85^Rev-Erb^ but not NHR-23^ROR^ (Fig. 4a). We mapped the regions of LIN-42^Period^ that are required for NHR-85^Rev-Erb^ binding and found that a minimal 51aa fragment present in both major LIN-42^Period^ isoforms is sufficient to mediate interactions (Fig. 5a). This domain is distinct from the interaction motifs implicated in mammalian Per2 and Rev-Erb ^26^. Because Per2 has been shown to interact with many mammalian NHRs, we extended our two-hybrid analysis of LIN-42^Period^ interactors to determine if LIN-42^Period^ also binds additional NHRs. We performed two-hybrid experiments between LIN-42^Period^ isoforms and 241 of the remaining 282 encoded *C. elegans* NHRs. We identified 65 NHRs that physically interact with LIN-42^Period^(Extended Data Fig. 5a,b). The additional interacting NHRs included DAF-12, which regulates the expression of the *let-7*-family of miRNAs and controls dauer development^28-30^, and NHR-14^HNF4a^, NHR-69^HNF4a^, and NHR-119 PPARα (Extended Data Fig. 5a,b) whose orthologs are also bound by Per2 ^26^. These findings suggest that many physical interactions between Period orthologs have been maintained since the divergence of nematodes and man and are, therefore, likely functional.

**Fig. 4.**
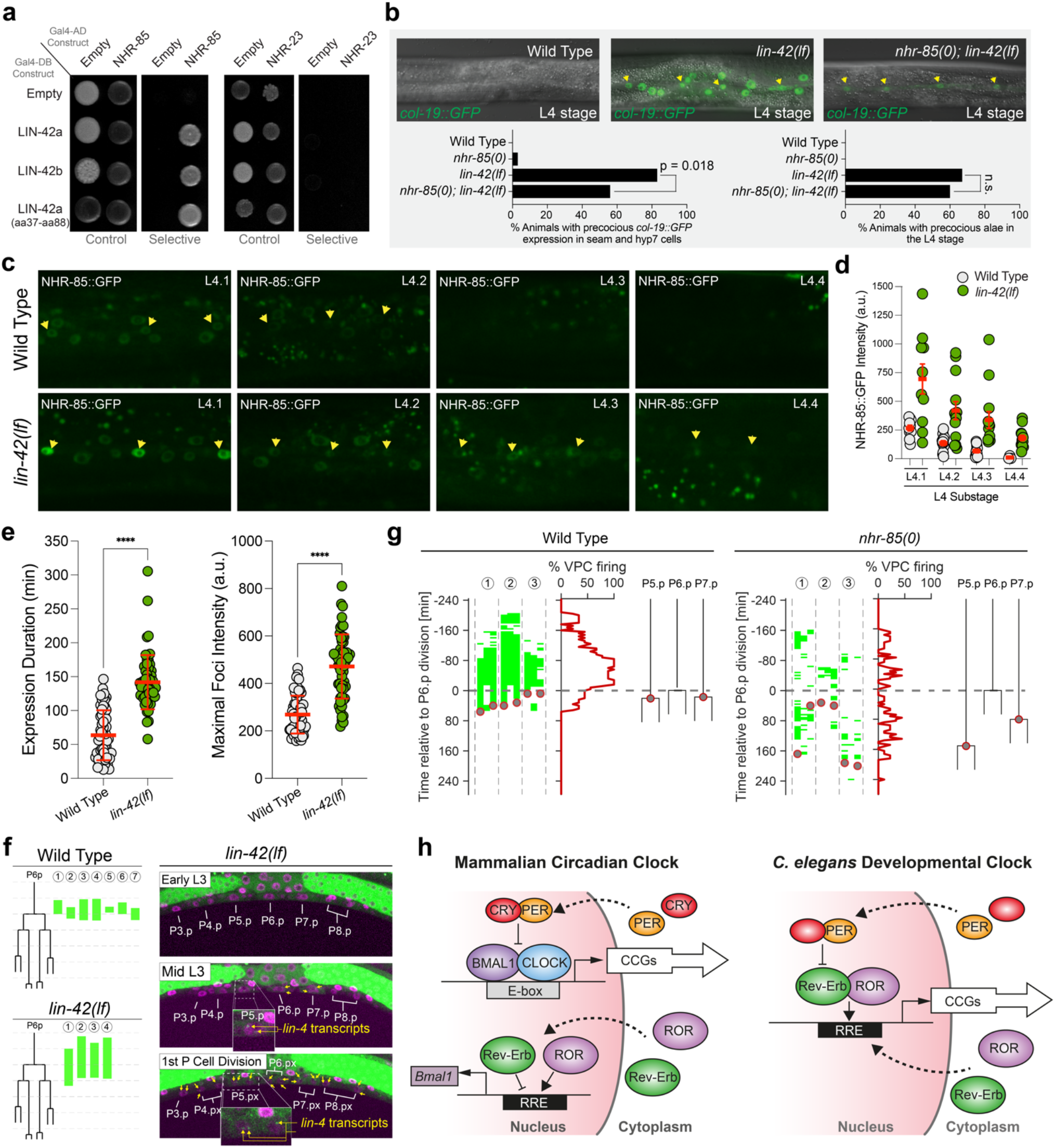
LIN-42^Period^ binds to and regulates the expression dynamics of NHR-85 to control the amplitude and duration of *lin-4* transcription. (**a**) LIN-42 isoforms interact with NHR-85^Rev-Erb^ but not NHR-23^ROR^ in two-hybrid assays. (**b**) *lin-42(lf)* mutants express *col-19::GFP* during the L3 stage of development and deletion of *nhr-85(0)* suppresses these phenotypes. Yellow arrows indicate the lateral seam cells of L4-staged animals. Error bars were calculated using two-tailed chi-square analysis. (**c and d**) Representative images and quantification of NHR-85^Rev-Erb^::GFP expression dynamics in hypodermal cells of L4-staged wild-type and *lin-4(lf)* animals. Yellow arrows indicate the lateral seam cells of L4-staged animals. Circles represent the average expression in three seam cells of an individual animal. Error bars show mean and standard deviation. Significance was calculated using Welch’s t-test. (**E**) Quantification of the duration and intensity of MCP-GFP foci in L3-staged animals. Error bars and significance are calculated as in d. (**f**) Time course analysis of the onset/offset times for MCP-GFP foci in VPCs of wild-type and *lin-42(lf)*. Green lines indicate the timing of *lin-4::24xMS2* expression in P6.p cells of individual animals. Pictographs show a representative image of the ventral surface of a single *lin-42(lf)* animal throughout the time course. (**g**) High-resolution time course analysis of *lin-4::24xMS2* expression in wild-type and *nhr-85(0)* mutants, aligned to first P6.p cell division (t=0). Green areas indicate detectable MCP-GFP foci in individual P cells (P5.p – P7.p) (n = 3 animals). Grey circles represent the timing of the P5.p and P7.p divisions. (**h**) Model of the mammalian circadian clock and the *C. elegans* developmental clock uncovered in this study.

Given the physical interaction between NHR-85^Rev-Erb^ and LIN-42^Period^, we asked whether NHR-85^Rev-Erb^ expression was required for the precocious phenotypes seen in *lin-42(lf)* mutants (*lin-42(n1089)*). We found that *lin-42(lf)* heterochronic phenotypes are partially ameliorated by removing *nhr-85* function. Specifically, the precocious expression of adult-specific reporters (e.g., *col-19::GFP)* in both seam and hyp7 cells observed in *lin-42(lf)* mutants is suppressed by *nhr-85* deletion; leaving weak expression in seam cells in double mutants, while precocious deposition of adult alae was not suppressed (Fig. 4b). To examine whether LIN-42^Period^ regulates NHR-85^Rev-Erb^ temporal expression, we compared the dynamics of NHR-85^Rev-Erb^::GFP (and NHR-23^ROR^::mScarlet) levels in wild-type and *lin-42(lf)* mutants. We found that the expression of NHR-85^Rev-Erb^ is altered in two ways by the *lin-42* mutation. First, the expression of NHR-85^Rev-Erb^::GFP is ∼2.3x more abundant at the beginning of the L4 stage in *lin-42* mutants when compared to wild-type animals (Fig. 4c,d). More importantly, the periodic dampening of NHR-85^Rev-Erb^ expression that usually occurs by the L4.2 stage of vulval morphogenesis (Fig. 3c,d) is altered in *lin-42(lf)* mutants. Specifically, NHR-85^Rev-Erb^::GFP expression perdures into the L4.4 stage *lin-42(lf)* animals (Fig. 4c,d). Mutations in *lin-42* do not alter the onset or duration of NHR-23^ROR^::mScarlet accumulation in hypodermal or vulval cells. This suggests that LIN-42 regulates *lin-4* transcriptional output by controlling the duration of NHR-85^Rev-Erb^/NHR-23^ROR^ heterodimeric complex formation in a manner that directly correlates with NHR-85 abundance.

Given the above hypothesis, we anticipated that specific features of *lin-4* transcriptional pulses would be altered in *lin-42* mutants. While most wild-type seam cells exhibit MCP-GFP foci during each larval stage (76%; n=66 nuclei), the percentage of seam cells showing detectable *lin-4* transcription is dramatically increased in *lin-42* mutants (100%; n=58 nuclei). This indicates that *lin-42* normally dampens *lin-4* transcriptional pulses in wild-type animals. In addition to elevating the likelihood that *lin-4::24xMS2* transcription is above a threshold sufficient to generate measurable MCP-GFP foci, *lin-42(lf)* mutations lead to an increase in the intensity of MCP-GFP foci indicating that LIN-42^Period^ normally also limits the rate of transcriptional activation of the *lin-4* locus (Fig. 4e). Time course experiments also revealed that overall duration of transcription events in lateral seam cells was ∼2.2 times longer in *lin-42* mutants compared to wild-type (Fig. 4e). Consistent with our hypothesis that the duration of *lin-4* transcription is dependent on the duration of NHR-85^Rev-Erb^ expression, the transcriptional onset of *lin-4::24xMS2* occurs earlier in L3 staged *lin-42* mutants and inappropriately extended through both the first and second Pn.p divisions (Fig. 4f).

While *nhr-85(0)* mutants did not misexpress adult-specific reporters (Fig. 5b), high-resolution imaging of VPC divisions and *lin-4::24xMS2* expression indicate two features of developmental timing were altered. In wild-type animals, transcriptional pulses of *lin-4::24xMS2* are both robust and concordant in adjacent wild-type VPCs (Figs. 4g & 1c). In contrast, MCP-GFP foci in *nhr-85(0)* mutants begin to accumulate at the same relative phase of L3-stage VPC development but are dimmer and only transiently observed (Fig. 4g). Second, under identical imaging conditions, the rapid and highly coordinated VPC divisions observed in wild-type animals is altered in *nhr-85(0)* mutants with some P5.p and P7.p dividing hours after the first P6.p division (Fig. 4g). These results indicate that NHR-85 functions to enhance the robustness of temporally regulated processes during development and that some level of *lin-4* transcription occurs without NHR-85^Rev-Erb^, perhaps driven by NHR-23^ROR^ alone. RNAi-mediated depletion of *nhr-23* activity in wild-type animals resulted in mild heterochronic phenotypes (Extended Data Fig. 6). Consistent with the hypothesis that NHR-23 and NHR-85 function cooperatively to control temporal regulation, the penetrance of these phenotypes was enhanced when *nhr-23* was also depleted in *nhr-85(0)* animals (Extended Data Fig. 6).

## Discussion

This study reveals the gene regulatory network controlling *lin-4* transcriptional pulses. This network shares integral components with the human circadian clock but exhibits essential differences in its regulatory architecture. In the circadian clock regulatory network, transcription factors CLOCK and BMAL1 generate rhythmic expression patterns of clock control genes (CCGs) ^31-34^ (Fig. 5h), including two core transcriptional repressors, Period and CRY as well as NHR genes Rev-Erb and ROR. Negative feedback on CLOCK/BMAL1 expression by Period/CRY heterodimers is essential for the proper generation of circadian rhythms. Rev-Erb and ROR modulate CLOCK and BMAL1 expression through opposing transcriptional activities (Fig. 4h) ^5-7^ but are dispensable for the generation of circadian oscillations ^35^. Instead of CLOCK and BMAL1, which are lacking in the *C. elegans* genome, we propose here that the worm orthologs of Rev-Erb and ROR compose the central transcription factors of the hypodermal developmental clock (Fig. 5h). In contrast to their antagonistic roles in the circadian clock, *C. elegans* NHR-85^Rev-Erb^ and NHR-23^ROR^ heterodimerize and work cooperatively to induce transcriptional pulses (Fig. 4h). The evolutionary rewiring of this interaction results in the phased expression of NHR-85^Rev-Erb^ and NHR-23^ROR^ generating a short temporal window for these proteins to cooperativity promote transcription. This enables transcription to be precisely dosed and timed at each larval stage. This rewiring also provides a direct physical link to the conserved opposing arm of the clock, LIN-42^Period^, which binds to NHR-85^Rev-Erb^ to control NHR-85^Rev-Erb^ expression post-transcriptionally (Fig. 4h). Additional physical and functional interactions between Period orthologs and conserved NHRs are maintained in the mammalian and *C. elegans* systems ^26,28^ and suggest that *C. elegans* NHRs, in addition to NHR-85^Rev-Erb^, may also function in temporal patterning.

While NHR-85^Rev-Erb^/NHR-23^ROR^ heterodimeric complexes generate pulses of *lin-4* transcription, NHR-23^ROR^ may play a more pervasive role in coordinating periodic transcription with overall animal development. Unlike *nhr-85, nhr-23* is an essential gene, and *nhr-23(0)* worms arrest during late embryogenesis/hatching, a time when oscillatory transcription begins^17,36,37^. Depletion of NHR-23^ROR^ during post-embryonic development results in highly penetrant larval arrest phenotypes ^38^. These arrests prevent somatic cell proliferation/differentiation, occur at the beginning of each larval stage, and correlate with the developmental periods where NHR-23^ROR^ expression peaks^38^. While the developmental arrest in NHR-23^ROR^-depleted animals makes it unfeasible to measure changes in transcription for individual genes, these arrests resemble post-embryonic developmental checkpoints where somatic cell proliferation and cyclical gene expression patterns (including *lin-4*) are halted during acute food removal/starvation ^11,39^. Starvation-induced checkpoints are mediated through the regulation of the conserved insulin signaling pathway, which generates various sterol-derived hormones ^40^. These hormones may serve as ligands for individual NHRs and provide a mechanism to organize global gene expression patterns. This type of coordination may underly one mechanism *C. elegans* larvae employ to generate identical developmental outcomes in diverse and rapidly changing environments.

