## Supplemental Figures + Materials and Methods for "A circadian-like gene network regulates heterochronic miRNA transcription in *C. elegans*"

*These authors contributed equally

**Acknowledgements:**

We would like to thank M. Walhout, J. Ward, J. Kimble for strains and reagents and D. Matus and T. Medwig-Kinney for help with confocal imaging. We would like to also thank G. Sumati for help with recombinant protein expression. L.J. is an investigator of the Howard Hughes Medical Institute. We acknowledge T. Medwig-Kinney, C. Vakoc, R. Martienssen, U. Pedmale, and D. Jackson for suggestions throughout the course of this study.

**Funding:**
National Institutes of Health R01GM117406 (CMH, NS, KH-M)
National Science Foundation 2217560 (CMH, JW)
CSHL Cancer Training Grant T32CA148056 (BK)
The CNRS ATIP/Avenir program Conseil Regional d'Île de France (DIM ELICIT-AAP-2020 KEIL) (20002719) (WK)
Fondation pour la Recherche Médicale FDT202204015083 (SS)
Howard Hughes Medical Institute (LJ, DA, MJ)

**Author Contributions:**
Conceptualization: CMH, BK, SS, WK
Methodology: BK, SS, NS, KH-M, DA, JW, MJ, LJ, WK, CMH
Investigation: BK, SS, DA, NS, KH-M, WK, CMH
Visualization: CMH, WK, BK, SS
Funding acquisition: CMH, WK, JT
Project administration: CMH, WK
Supervision: CMH, WK, JT
Writing – original draft: CMH
Writing – review & editing: CMH, WK, BK

**Competing interests:** All authors declare no competing interests.

**Data and materials availability:** Strains and plasmids listed in the Supplementary Materials and Methods Section are available through the *Caenorhabditis Genetics* Center or by request to C.M.H.. The authors affirm that all data necessary for confirming the conclusions of the article are present within the article, figures, and tables.

**Supplementary Materials**

Materials and Methods

Extended Data Figs. S1-S6

Tables S1-S4

References (*1-20*)

Movies S1-S2

**SUPPLEMENTAL FIGURES:**

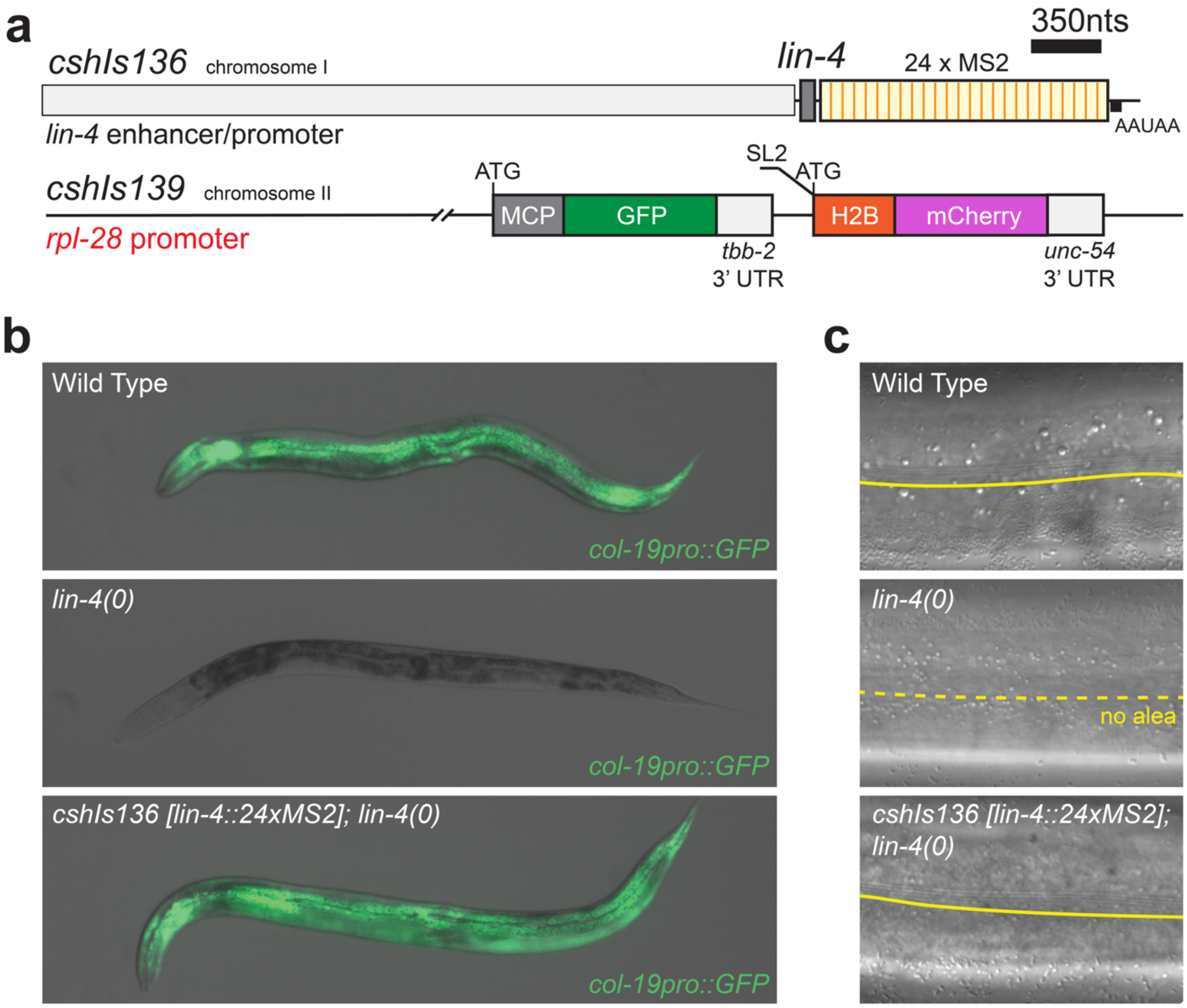


**Extended Data Fig. S1. The single copy, MS2-tagged *lin-4* transgene rescues *lin-4* temporal patterning defects.** (**a**) I diagram illustrating the two transgenes utilized in the lin-4::24xMS2/MCP-GFP system. (**b**) Wild-type animals express a *col-19::GFP* transgene immediately after the L4 molt (100%; n = 50) while *lin-4(e912)* mutants fail to induce *col-19::GFP* expression (0% expressing; n = 43). *cshIs136 [lin-4::24xMS2; unc-119(+) I; lin-4(e912) I]* animals exhibit normal *col-19::GFP* expression (100% expressing; n = 30). (**c**) Wild-type animals generate adult-specific alae structures on the cuticles of young adult animals (100%; n = 23). 100% (n = 40) of *lin-4(e912)* animals lack these structures. *cshIs136 [lin-4::24xMS2; unc-119(+) I; lin-4(e912) I*] animals exhibit wild-type adult alae (100%; n = 40).

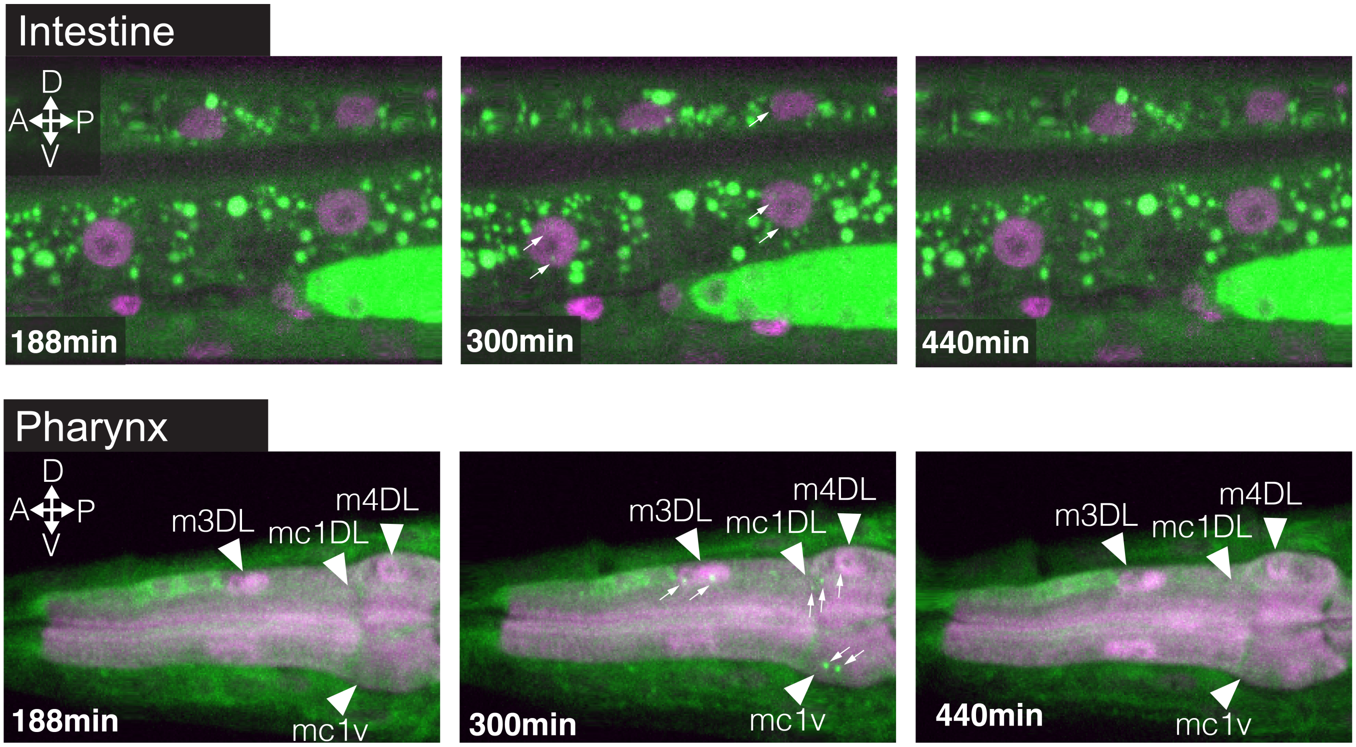


**Extended Data Fig. S2. *lin-4::24xMS2* expression is pulsatile in other somatic cells.** Timeseries of *lin-4::24xMS2* expression in intestine cells and cells of the pharynx in L3-staged animals. Small arrows indicate MCP-GFP foci and triangles indicate the location of the indicated cell.

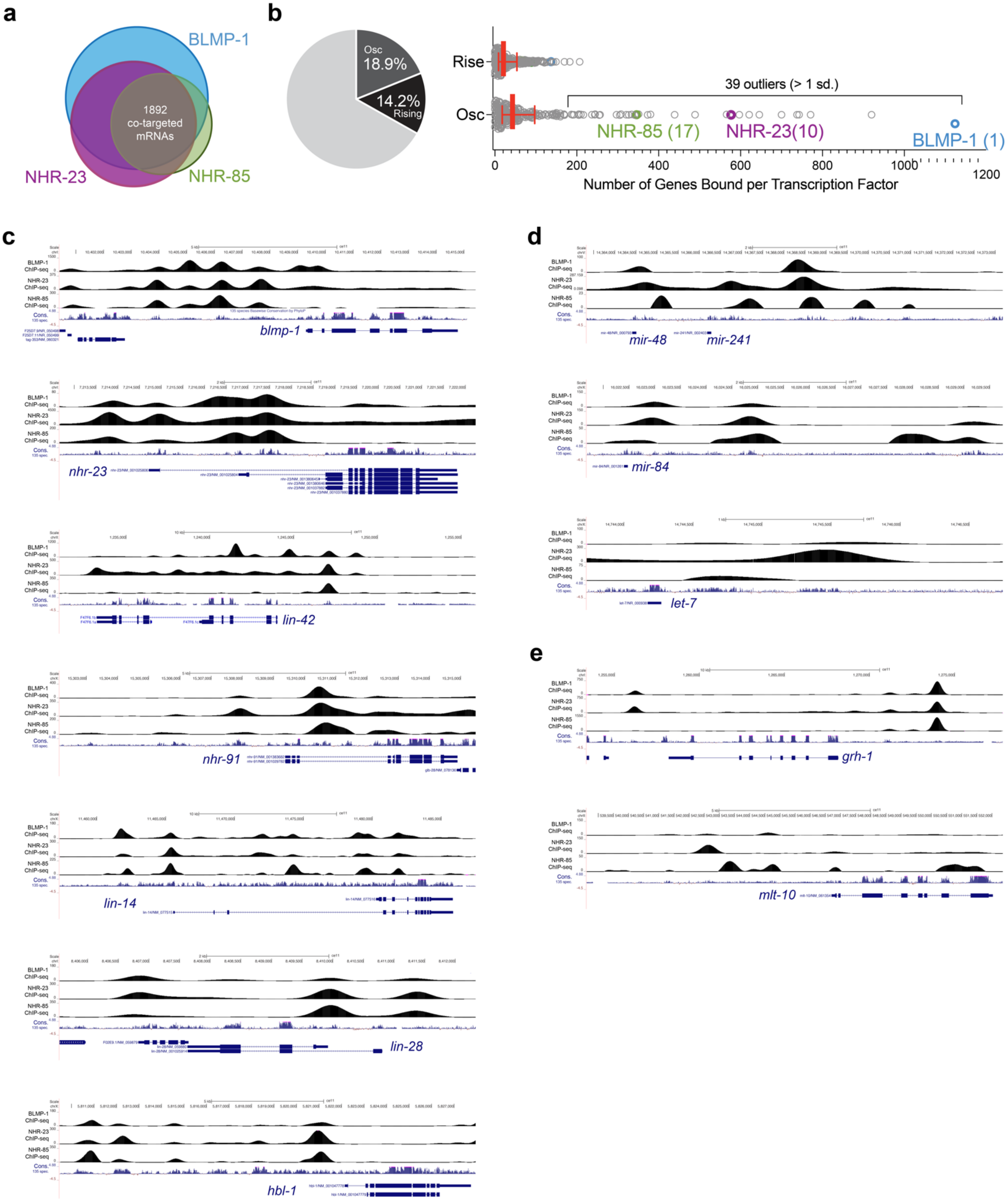
a
BLMP-1
1892
co-targeted
mRNAs
NHR-23
NHR-85
b
Osc
18.9%
14.2%
Rising
Rise
Osc
39 outliers (> 1 sd.)
NHR-85 (17)
NHR-23(10)
BLMP-1 (1)
0
200
400
600
800
1000
1200
Number of Genes Bound per Transcription Factor
c
BLMP-1 ChIP-seq
NHR-23 ChIP-seq
NHR-85 ChIP-seq
Cons.
blmp-1
nhr-23
lin-42
nhr-91
lin-14
lin-28
hbl-1
d
mir-48
mir-241
mir-84
let-7
e
grh-1
mlt-10

**Extended Data Fig. S3. BLMP-1, NHR-23$^{ROR}$, and NHR-85$^{Rev\text{-}Erb}$ binding sites are enriched in the promoters of cyclically transcribed genes as well as genes that function in the heterochronic pathway.** (**a**) Venn diagram indicating the level of overlap of genes targeted by BLMP-1, NHR-85 and NHR-23. For this analysis, peaks from NHR-85$^{Rev\text{-}Erb}$, NHR-23$^{ROR}$ and BLMP-1 ChIP-seq experiments were assigned to the nearest proximal protein coding genes (+3kb upstream and 300bp downstream) (Tables S1 and S2). (**b**) NHR-85$^{Rev\text{-}Erb}$, NHR-23$^{ROR}$ and BLMP-1 binding sites are enriched in the promoters of genes that exhibit oscillatory expression patterns. Calculations were derived from population-based RNA-seq time course data sets (*1*) that partition the expression of post-embryonically expressed genes into three types: (i) transcripts with their transcriptional periodicity tied to the molting cycle (osc; 18.9%; 2,718 of 14,378 total genes) (ii) genes with rising expression throughout larval development (rising; 14.2%) and (iii) genes with flat expression profile (flat; 66.9%) (Fig. 2E). Using these data sets, we compared the numbers of cyclically expressed genes that were targeted by NHR-85$^{Rev\text{-}Erb}$ or NHR-23$^{ROR}$ to those targeted by any of the other 170 TFs queried in by the *C. elegans* ModEncode Project (using 265 publicly available, validated ModEncode ChIP-seq data-sets (tables S1 and S2)(*2*). (**c**) Browser tracks overlaying the BLMP-1, NHR-23$^{ROR}$, and NHR-85$^{Rev\text{-}Erb}$ ChIP-seq signal near the indicated loci.

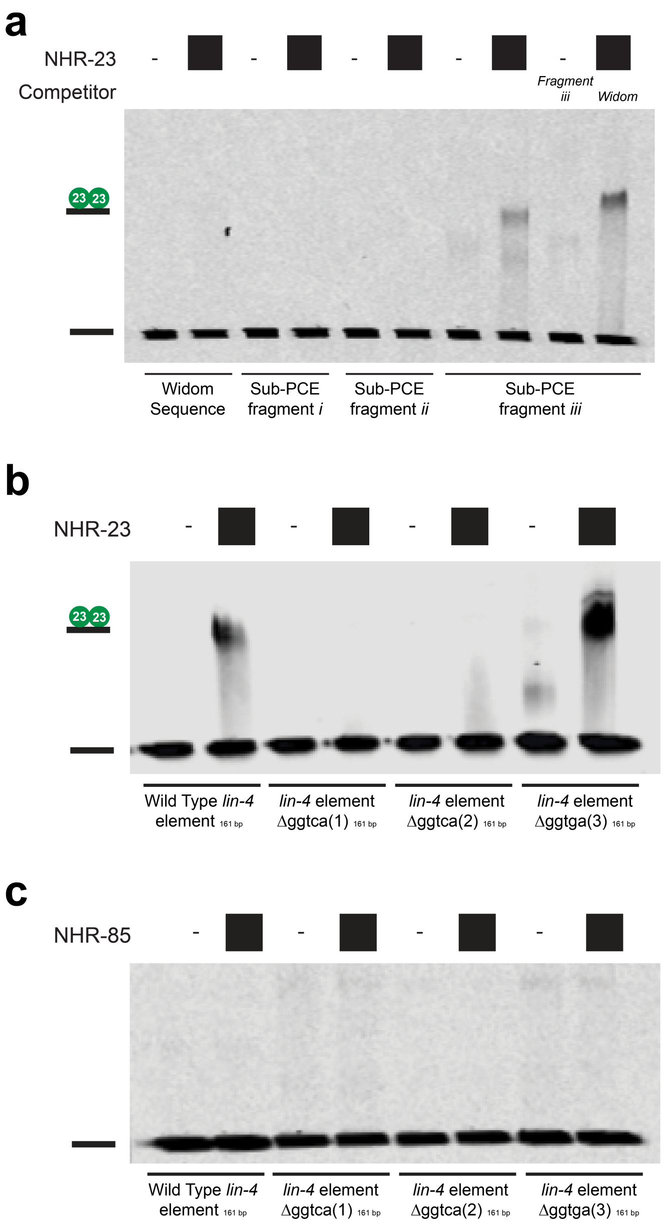


**Extended Data Fig. S4. NHR-23$^{ROR}$ can bind the *lin-4* sub-fragment III at high protein concentrations while NHR-85$^{Rev\text{-}Erb}$ cannot.** (**a**) NHR-23$^{ROR}$ can bind specifically to PCE sub-fragment III that contains both direct repeats outlined in Fig. 2F and G. (**b**) NHR-23$^{ROR}$ binding to the PCE sub-fragment III requires the two direct repeats but does not require the un-conserved single GGTGA element downstream of the direct repeats (see Fig. 2 for details). (**c**) NHR-85$^{Rev\text{-}Erb}$ does not bind to any of the PCE sub-elements to any appreciable degree.

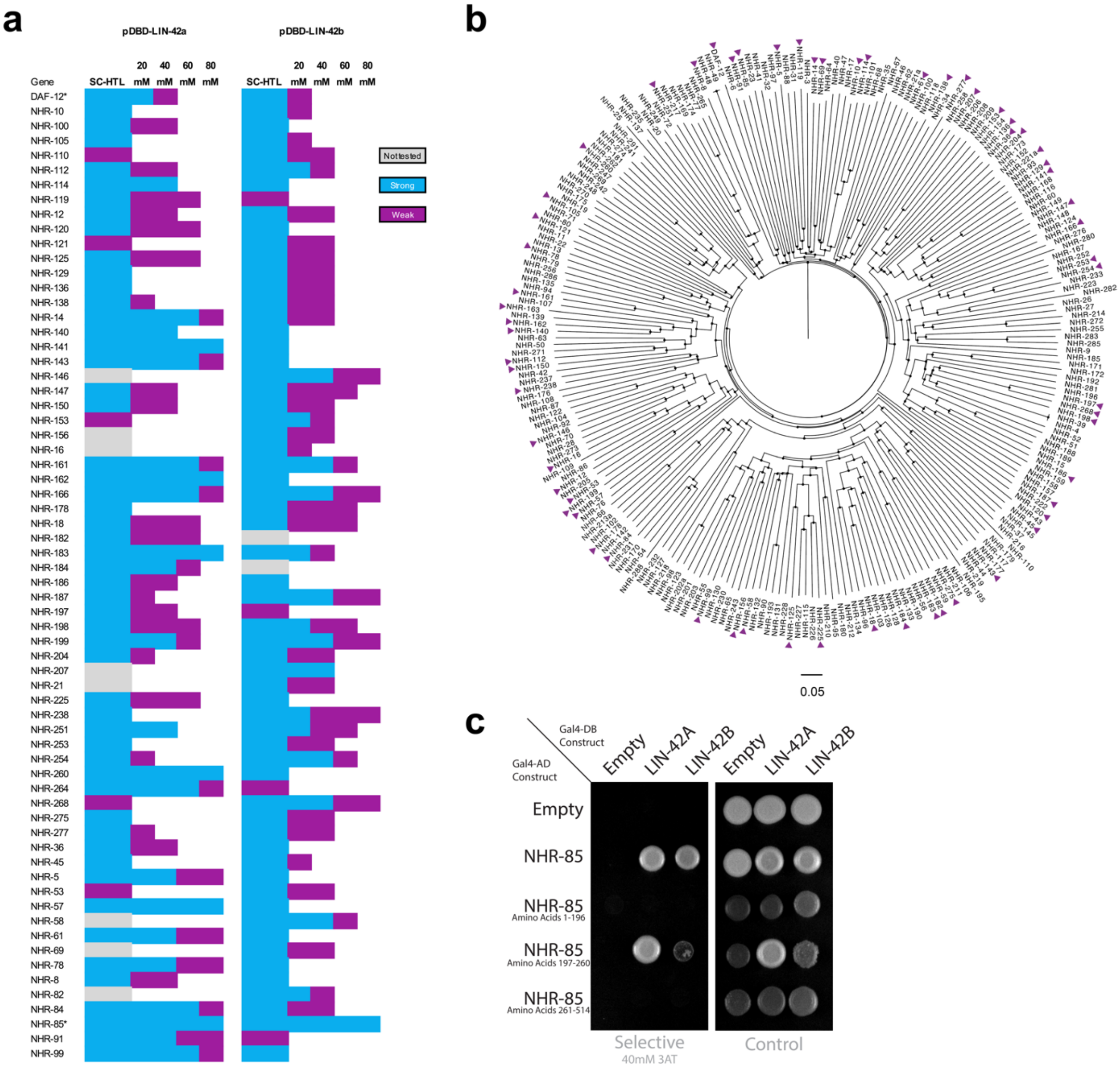


**Extended Data Fig. S5. LIN-42$^{Period}$ interacts with multiple C. elegans NHRs and interacts with NHR-85$^{Rev\text{-}Erb}$ via the hinge region that separates the DNA-binding domain and ligand-binding domain.** (**a**) A list depicting the NHRs that interact with one of the two two-hybrid baits for LIN-42 (LIN-42a and LIN-42b isoforms). Colors indicate the level of yeast growth solid growth media containing the indicated concentration of 3-AT. (**b**) Dendrogram of *C. elegans* nuclear hormone receptors showing the relationships between conserved sub-families of NHRs and the distribution of LIN-42 interactions within this large family of TFs. Purple triangles indicate NHRs that interact with one or both LIN-42 isoforms. (**c**) Two hybrid results that indicate that both LIN-42 isoforms can interact with the central hinge region of NHR-85.

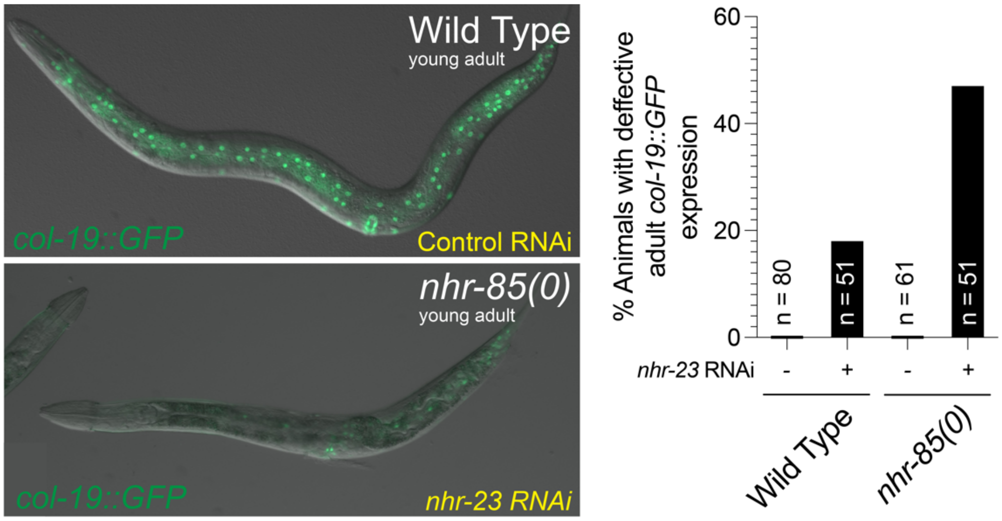


**Fig. S6. Depletion of nhr-23 results in defects in expressing adult-specific GFP reporters after the L4 molt.** The adult-specific *col-19::GFP* reporter is expressed in both seam and hyp7 cells in wild-type animals. Depletion of *nhr-23* prevents normal expression of *col-19::GFP* during adulthood. In *nhr-23* RNAi treated animals, the indicated percentage of animals expressed *col-19::GFP* in the head and tail cells of the hypodermis. The reduction of *col-19::GFP* expression in young adult animals is stronger in animals lacking *nhr-85*. *nhr-23* RNAi cultures were diluted to 50% with control RNAi bacteria to enable animals to develop to adulthood and avoid the fully penetrant larval arrest phenotypes associated with undiluted *nhr-23* RNAi cultures.

**MATERIALS AND METHODS:**

***C. elegans* maintenance and genetics.**

*C. elegans* strains were maintained on standard media at 20°C and fed *E. coli* OP50 (*3*). Some strains were provided by the CGC, which is funded by NIH Office of Research Infrastructure Programs (P40 OD010440). A complete list of strains is outlined in Table S3.

**RNAi Feeding.**

RNAi by feeding was performed using *E. coli* HT115 expressing double stranded RNA corresponding to the indicated target gene or containing a control dsRNA expression plasmid that does not contain sequences corresponding to any *C. elegans* gene (*4, 5*). To prevent contamination by *E. coli* OP50, L4-staged animals were added to RNAi plates individually after removing co-transferred bacteria. For RNAi against *nhr-23*, bacterial cultures were diluted with control RNAi cultures at the indicated levels prior to experimental onset. In experiments in Fig. S6, starved L1 animals of the indicated genotypes were used. Unless otherwise noted, F1 progeny were analyzed for RNAi-induced phenotypes in the following 2-4 days. Plasmids used for RNAi are outlined in Table S4.

**CRISPR genome editing.**

Genome editing and transgene insertion was accomplished using standard CRISPR/Cas9-mediated genomic editing to the ttTi5605 or ttTi4348 landing site following standard protocols (*6*). For CRISPR/Cas9 editing of the endogenous *lin-42* gene, pCMH1434 (expressing Cas9 and a synthetic CRISPR guide RNA targeting a genomic region encoding the LIN-42 C-terminus) and pCMH1439 (encoding a LIN-42::YFP fragment) were injected in to N2 animals and screened by PCR to identify transgene insertion at the *lin-42* gene. Plasmids used to generate CRISPR alleles are outlined in Table S4.

**Microfluidics and long-term imaging.**

For microfluidics experiments, early to mid L1-staged animals were isolated 6h after starvation-induced L1 arrest at 20°C before an experimental time course. Other stages were individually isolated by observing defined cellular and morphological features indicative of animals developmental stage (*7*). Animals were mounted into the microfluidic device as previously described (*8*). During the imaging process, animals were constantly fed NA22 *E. coli* suspended in S medium. The temperature was kept constant at 20˚C both, at the objective and the microfluidic device using a custom-built water-cooled aluminum ring (for objective) and custom-built aluminum stage inset that was directly coupled to a thermal Peltier device.

**Image acquisition**

MS2-MCP-GFP live imaging Live imaging was performed with a 60x, 1.2NA objective on a Nikon Ti2 Eclipse microscope equipped with a V3 CREST spinning disk confocal module. To ensure fast multichannel acquisition, hardware triggering was implemented between a MadCityLabs NANO Z200-N piezo z-stage, a Photometrics Prime 95B sCMOS camera with 25mm field of view (2048x2048 pixels, pixel size 11um corresponding to 183nm), and a Lumencor© Celesta solid-state laser source via a National Instruments (NI) PCIe-6323 card. Laser wavelengths of 488nm and 545nm were used to excite MCP-GFP and histone-mCherry, respectively. Acquiring a dual-color z-stack with 51 slices and 50ms exposure times takes approximately 3.2 seconds with this setup.

Confocal Microscopy: Images were acquired using a Hamamatsu Orca EM-CCD camera and a Borealis-modified Yokagawa CSU-10 spinning disk confocal microscope (Nobska Imaging, Inc.) with a Plan-APOCHROMAT x 100/1.4 or 40/1.4 oil DIC objective controlled by MetaMorph software (version: 7.8.12.0). Animals were anesthetized on 5% agarose pads containing 10mM sodium azide and secured with a coverslip. Imaging on the microfluidic device was performed on a Zeiss AXIO Observer.Z7 inverted microscope using a 40X glycerol immersion objective and DIC and GFP filters controlled by ZEN software (version 2.5). Images were captured using a Hamamatsu C11440 digital camera. For scoring plate level phenotypes, images were acquired using a Moticam CMOS (Motic) camera attached to a Zeiss dissecting microscope.

Wide-field Fluorescence microscopy:

Images were acquired with a Zeiss Axio Observer microscope equipped with Nomarski and fluorescence optics as well as a Hamamatsu Orca Flash 4.0 FL Plus camera. An LED lamp emitting at 470 nm was used for fluorophore excitation. For single images, animals were immobilized on 2% agarose pads supplemented with 100mM Levamisole (Sigma). For single images, animals were immobilized on 2% agarose pads supplemented with 100mM Levamisole (Sigma). For long-term imaging methods, see Microfluidics and Long-term Imaging section.

**Fluorescent Reporter Quantification.**

Reporter lines were imaged using wide-field fluorescence or confocal microscopy as described above. The average intensity (arbitrary unites) per seam cell was measured using ImageJ. Each seam cell intensity was determined by the measurement of fluorescent intensity of the nucleus minus the intensity of a background sample. The fluorescent intensity of each animal was determined by the average of three seam cells. Ten animals per developmental stage were imaged unless otherwise noted.

**MCP-GFP live imaging**.

Long term imaging: For long-term live imaging across several larval stages, animals were mounted in a microfluidic chamber 6h after L1 arrest and grown on NA22 bacteria suspended in S-medium until mid L4 as previously described (*8*). At each time point, animals were *reversibly* immobilized using microfluidic pressures and flows. A z-stack of 51 images separated by 0.5um was acquired at four overlapping positions, covering the entire microfluidic chamber. Thereafter, the animal was released from immobilization and left to roam and feed freely until the next time point.

Short term imaging: For short-term live imaging (Figs. 1, 4 & 5) developmentally staged animals were mounted into the microfluidic chamber as previously described (*8*), a few hours before the expected appearance of MS2 spots. Minutes before the appearance of MS2 spots, animals were immobilized using microfluidic pressures and flows and kept immobilized for the entire duration of the experiment. This allowed to maintain a stable worm position enabling automated analyses (see below). At each time point (every 4min), a stack of 21 images separated by 0.5um was acquired at four overlapping positions, covering the entire microfluidic chamber. Occasionally, animals arrested development upon prolonged immobilizations, as evident by the cessation of germ line divisions (L2-L4 larvae), the cell-cycle arrest of vulval precursor cells (VCPs) (L3 larvae) or by failure to advance through vulval morphogenesis (L4 larvae). These animals were excluded from further analysis. As opposed to all other genotypes imaged, *nhr-85(0)* mutants animals exhibited a pronounced tendency to roll under these imaging conditions, precluding MS2 spot tracking within nuclei of the lateral hypodermis.

**MCP-GFP live imaging data analysis.**

Short term imaging:

All events (cell divisions, onset and offset of MCP-GFP spots) were scored manually in the time series. For short-term live imaging, all analysis was performed using custom-written FIJI macros, pixel classification with random forest trees in Ilastik and MATLAB© scripts. The main challenge in this analysis is residual animal movement between time points as well as low signal-to-noise ratio of the MCP-GFP signal.

Long term imaging and 3D tracking:

First, the worm backbone was detected in each frame by skeletonization, using a thresholded probability map, obtained by processing a maximum z-projection of the MCP-GFP channel, through a custom-trained Ilastik pixel classifier. Next, computational straightening was performed along this backbone (*8*) to obtain a time series of straightened worm z-stacks. This straightened time series has the advantage that residual movement is primarily along the anteroposterior animal axis. Next, we divided the straightened worm z-stack along the worm axis into overlapping (20%) segments of 500 pixels (~91um) in length. Each of these segments was then manually registered to obtain z-stacks in which tracking of almost all hypodermal nuclei could automatically performed with minimal user corrections. To improve nuclear signal for segmentation, the histone-mCherry signal was augmented, using a custom-trained Ilastik pixel classifier. 3D tracking and manual correction was then performed on the resulting Ilastik probability maps using the FIJI TrackMate plugin (*9*) with LoG detector and LAP tracker. For each segment, frames with substantial animal movement *between* z-slices were excluded from subsequent analysis. Whenever nuclei were tracked twice (due to overlapping worm segments), the nucleus with the most tracked time points was chosen for subsequent MCP-GFP spot analysis.

**MCP-GFP spot tracking and intensity quantification.**

MS2 spots were tracked in each tracked nucleus, using the FIJI TrackMate plugin with LoG detector and LAP tracker and the Ilastik nucleus probability maps as mask to include only spots inside nuclei. Each MCP-GFP spot track was manually corrected, and spots were added to frames occasionally, in which the TrackMate LoG detector failed to detect them. For quantification of MCP-GFP spot intensities, a 2D Gaussian fit to the maximum z-projection of three z-slices around the peak slice determines the position of the spot. Background was calculated as the average intensity in a ring between 3 and 5 pixels away from the spot position. The spot intensity was calculated by integrating the fluorescence over a circle with a radius of 2 pixels around the spot position and subtracting the estimated background.

**MS2-MCP-GFP trace analysis.**

MCP-GFP traces were smoothed with a Gaussian of width 1.5 frames (6 minutes). Using this filtered trace, for each tracked locus, duration and maximum intensity during the transcriptional pulse was determined. Loci were considered “ON”, if spot intensity was above 150 counts and “OFF” if below. Duration of transcription for each locus was determined as the time interval between the first “ON” time point and the last. Typically, loci stayed “ON” for the entire duration of the transcriptional pulse within a given larval stage.

**Yeast two-hybrid.**

Plasmids containing target proteins fused to GAL4 DNA-binding-domain (pDBD) and GAL-4 Activation Domain (pAD) were co-transformed into the pJ69-4a Y2H yeast strain (*10*) as previously described (*11, 12*). Transformed yeast was plated on SC-TRP-LEU plates for three days. Three colonies from each transformation plate were streaked onto SC-HIS-TRP-LEU plates containing 3-AT at the indicted concentrations. Protein interactions were determined by visible growth on 3-AT conditions with negative growth in empty vector controls after three days. For the large-scale LIN-42 screen, pDBD containing LIN-42a, LIN-42b, and the empty vector control were individually mated to each pAD construct from the WTF2.2 yeast library (Reece-Hoyes et al., 2011) (*13*). For visualization of results, individual colonies were grown overnight in YPD in 96-well plates. Overnight cultures were diluted 1/200 in ddH20 and 3μL was pipetted on to selective 3-AT and control plates. After three days of growth, plates were imaged on a Fotodyne FOTO/Analyst Investigator/FX darkroom imaging station. Plasmids used in 1- and 2-hybrid experiments are outlined in Table S4.

**Bioinformatic analysis of BLMP-1, NHR-82, and NHR-23 ChIP-seq data.**

ChIP-seq short reads were first clipped off adaptor sequences. Reads of minimum 22bp were mapped to the UCSC *C. elegans* genome (ce10) using bowtie program76 (*14*) looking for unique alignments with no more than 2 mismatches. MACS program (v1.4)(*15*) was used for peak calling with significant p-value cutoff equals 1e-5. Target annotations were based on WormBase (version 220) using customized R scripts and Bioconductor packages. ModeEncode data sets (*2*) (BLMP-1(ENCFF108AEB and ENCFF615AMZ), NHR-23(ENCFF019QMH) and NHR-85(ENCFF018KHA)) were used in the initial analysis and we defined the promoter region as upstream 3kbp to downstream 300bp around the transcription start site (TSS). Peaks located in promoter regions were annotated to their closest TSS sites for both coding and non-coding genes (Table S1). These potential targets were then overlapped to sets of oscillatory genes identified in a previous mRNA-seq-based study (*1*). Overlapping target gene sets organized in Figure 2D are from postembryonic ChIP-seq samples for each TF. Raw data for each of the 265 ModEncode data sets (annotation numbers listed individually in Table S1) used in Figure 2E were downloaded from modENCODE (*2*) and processed in the same way as BLMP-1, NHR-23 and NHR-85 datasets (*16*). ModEncode-derived peaks from each TF ChIP-seq dataset were compared to identify common sites with at least one base pair overlapping using BEDTools. All ChIP-seq mapping graphs and images were produced in R by customized scripts.

**Protein preparation.** Full-length *C. elegans* protein NHR-23 was cloned as a N-terminal Strep-SUMO fusion protein in a pFL vector of the MultiBac Baculovirus expression system to create pCMH1662 (*17*). This construct was expressed in insect *Sf9* cells grown in CCM3 media (HyClone) at 27°C for 60 h. Cells were harvested by spinning at 2200 rpm for 20 min and resuspended in lysis/wash buffer (20 mM Tris pH 8.0, 200 mM NaCl, 5 mM BME) and a protease inhibitor cocktail before flash freezing in liquid $N_2$. Cell pellets were stored at -80°C. Cell pellets were thawed and sonicated once. Poly-ethylene imine (PEI) was added at 0.2% to the lysate after cell pellets were thawed and sonicated. The lysate was then spun by ultracentrifuge at 38,000 rpm for 45 min, at 4°C. The supernatant of the lysate was then used for batch binding with Strep-Tactin superflow resin (IBA) for 1 hour while on a rolling shaker at 4°C. The affinity beads were harvested by spinning at 1000 rcf for 5 minutes, then resuspended in lysis/wash buffer and applied to a gravity column. The column was washed with 30 column volumes of lysis/wash buffer and 5 column volumes of ATP wash buffer (20 mM Tris pH 8.0, 200 mM NaCl, 5 mM BME, 2 mM ATP). The protein was eluted from the affinity column in 2 column volumes of elution buffer (20 mM Tris pH 8.0, 200 mM NaCl, 5 mM BME, 2 mM desthiobiotin). The Strep-SUMO tag was cleaved from NHR23 by ULP1* protease overnight at 4°C. The protein was then concentrated and loaded onto a 10/300 Superdex200 Increase gel filtration column (Cytiva Life Sciences), running in lysis/wash buffer, chromatographed for ~30 mL at 0.6 mL $min^{-1}$. Protein purity and cleavage efficiency was assessed by SDS-PAGE. Plasmids used to express recombinant proteins are outlined in Table S4.

Full length *C. elegans* protein NHR-85 was cloned as a N-terminal Strep-fusion protein in a pFL vector of the MultiBac Baculovirus expression system to create pCMH2206. NHR-85 was purified using the same method as above, with the exception of the overnight N-terminal tag cleavage step.

**Microscale thermophoresis analysis.** Binding assays of purified NHR23 or strep-NHR85 was measured using a Monolith NT.115 Pico running MO Control version 1.6 (NanoTemper Technologies). Assays were performed in 100 mM NaCl, 20 mM Tris pH 8.0, 0.05% Tween-20. AlexaFluor647 NHS Ester (ThermoFisher Scientific) labeled NHR-23 (200 pM) was mixed with 16 serial dilutions of strep-NHR-85 starting at 31.5 uM and loaded into microscale thermophoresis premium coated capillaries (NanoTemper Technologies). MST measurements were recorded at 25°C using 30% excitation power and 60% MST power. Measurements were performed in duplicate. Determination of the binding constant was performed using MO Affinity Analysis v.2.3.

AlexaFluor647 NHS Ester (ThermoFisher Scientific) labeled strep-NHR-85 (400 pM) was mixed with 16 serial dilutions of NHR-23 starting at 625 nM. MST measurements were recorded at 25°C using 15% excitation power and 40% MST power. Measurements were performed in triplicate. Determination of the binding constant was performed using MO Affinity Analysis v.2.3.

**Gel Shifts.** For gel shifts with free DNA, 5' IRDye (IRDye700 or IRDye800)-labeled and unlabeled oligos were obtained from IDT (Coralville, Iowa) and used to amplify DNA probes of the indicated sequences. For wild-type probes, the indicated PCE fragments were amplified pCMH1954. For mutants probes that harbor mutations in either of the GGTCA repeats, synthetic DNAs were obtained from Synbio Technologies (Manmouth Junction, NJ, USA) and used to amplify the corresponding mutant DNA fragments with the following sequences (mutations underlined and italicized):

C_ROR(1)REV:TTTGCATCCTCATTCTCAACACCTCGTTTTTTTCCCTTTTCTTGCACAAATTGA*cctgt*G
TCGGTCAGTAAACCCCCCCCCCCCCCCCCCCATTGAGGTGACCAATTGGTTTTTCTTTTCCTTTACT
TTCTCCTTCACTTTCTCTCTC TCGGATCACCAGC
C_REV:TTTGCATCCTCATTCTCAACACCTCGTTTTTTTCCCTTTTCTTGCACAAATTGAGGTCAGTC*c*
*ctgtc*TAAACCCCCCCCCCCCCCCCCCCATTGAGGTGACCAATTGGTTTTTCTTTTCCTTTACTTTCTC
CTTCACTTTCTCTCTC TCGGATCACCAGC
For binding reactions, recombinant proteins were incubated with gel purified DNA probes in 10 mM Tris pH 7.5, 50 mM KCl, 1 mM DTT, 0.1mg/mL poly (dIdC) (Sigma-Aldrich), and 0.25% Tween 20 for 30 minutes at $20^{\circ}$C (in dark chamber). Samples were then run in a 4% native polyacrylamide gel containing 50mM Tris pH 7.5, 0.38 M glycine and 2mM EDTA in 1x TBE buffer. Gels were imaged and quantified using a Li-Cor Odyssey Imager (Lincoln, Nebraska).

**Graph Plots and Statistical Analysis.** Plots and diagrams were generated using GraphPad Prism v9 (GraphPad Software, San Diego, Ca) or custom-written MATLAB® scripts. Statistical significance was determined using a two-tailed un-paired Student's t-test. P<0.05 was considered statistically significant. **** indicates P < 0.0001.

Table S1. Annotation of ChiP-seq Data from ModEncode used in Fig. S3.
Table S2. Table depicting the distribution of ChIP-seq target genes for each TF parsed out into expression categories outlined in Fig. S3.
Table S3. *C. elegans* and yeast strains used this work.
Table S4. List of plasmids used in this work.
